# Integrated bioinformatics and wet-lab analysis revealed prominent inflammatory genes of Extracellular Matrix as prognostic biomarkers in patients with advance IBD requiring early surgery

**DOI:** 10.1101/2023.12.13.570869

**Authors:** Farzad Dehghani Mahmoudabadi, Binazir Khanabadi

**Affiliations:** Department of General Surgery, Medical Science of Shahid Beheshti University, Tehran, Iran; Basic and Molecular Epidemiology of Gastrointestinal Disorders Research Center, Research Institute for Gastroenterology and Liver Diseases, Shahid Beheshti University of Medical Sciences, Tehran, Iran

**Keywords:** Advance IBD, Early surgery, prognostic biomarkers, ECM

## Abstract

**Background:** The number of patients with inflammatory bowel disease (IBD) is increasing worldwide. Due to the fact that at the age of 20 to 30 years, this autoimmune disease is very common; Investigating and identifying prognostic biomarkers in advanced IBD is very important; Because according to the identification of these biomarkers, patients who need early surgery can be nominated and undergo surgery without wasting time and treatment costs. In this study, with the aim of identifying effective biomarkers involved in the inflammatory part of extracellular matrix (ECM) in the early surgery of IBD, separately from Crohns disease and ulcerative colitis.

**Method:** In this study, we examined 50 patients in both patient groups as well as the normal group. The expression of the nominated genes MASP2, DKC1, HNF4A, and STAT3 was analyzed using quantitative polymerase chain reaction (Q-PCR) and relative quantification was determined using the 2^-ΔΔCt^ method. ROC curve analysis was performed to compare IBD (UC & CD) and normal for the investigated genes. The correlation between adhesion molecule gene expression and immunophenotype was analyzed. Also we comprehensively analyzed the genetic alteration, prognostic value and gene regulatory networks using multiple databases.

**Result:** The obtained results showed that MASP2 and DKC1 genes were significantly expressed in advanced UC patients, as well as HNF4A and STAT3 in advanced CD patients.

**Conclusion:** It can be stated that the biomarker panel MASP2, DKC1, HNF4A, and STAT3 related to them have a significant prognostic role in the candidates of IBD patients for early surgery.

**Graphical abstract:** 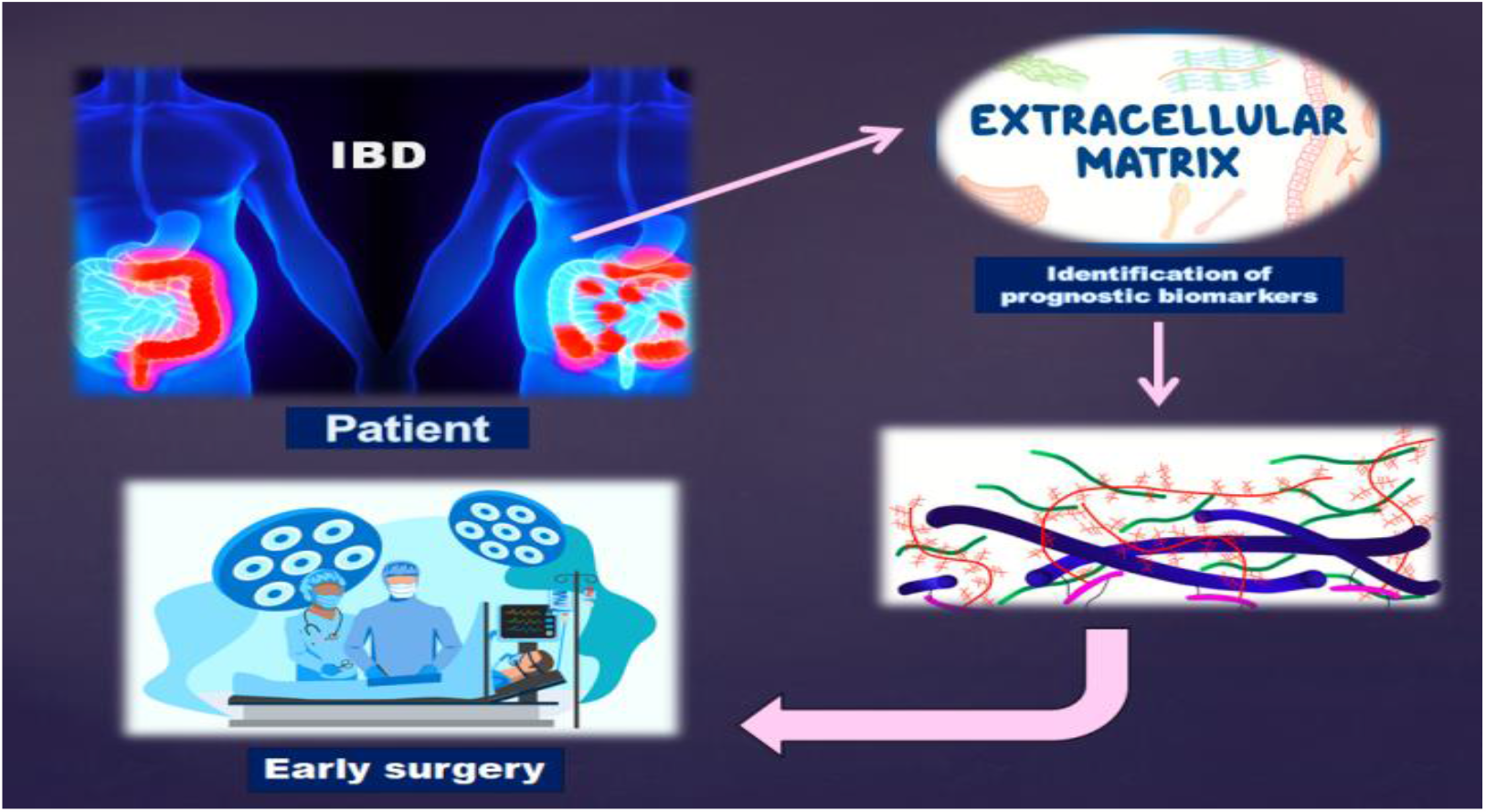

## 1. Introduction

Inflammatory bowel disease (IBD) (Crohns and colitis) is an idiopathic disorder in the digestive system that is characterized by chronic and segmental inflammation and can usually be recurrent(1, 2). IBD also leads to an increased risk of colorectal cancer (CRC) that is up to 30 times higher than in the general population (3). The wide variety of intestinal symptoms and extra intestinal manifestations, with different phenotypes of disease behavior, make the use of &treatment to target; more challenging(4).In the last two decades, the emergence of methods biological therapy and modification of therapeutic paradigm have revolutionized the medical management of IBD(5). Specific and significant improvements include: the introduction of tumor necrosis factor (TNF) antagonists and integrin inhibitors, the use of combination treatments, the monitoring of therapeutic drugs and the initiation of effective treatments in high-risk patients(6, 7); But despite all these fruitful points, one of the biggest challenges in the implementation of these strategies are two very important points: a. Determining which patients are more at risk of complicated complications of IBD and how long drug protocols should be continued in these people; b. Also, when is the most appropriate time for surgery in these patients (7). The action to make these decisions is usually based on the judgment of the patients; clinical manifestations, and because of the patients activities that are evaluated by the symptoms, it can have serious consequences; Based on this, identifying the patients who have the highest risk of complications and disease progression, as well as benefiting these people from determining the appropriate time for effective treatment or surgery, will be a valuable goal(6, 8). In the development of IBD and Inflammatory tissues, various cellular and molecular functions are involved, transcription, cell cycle control, cell adhesion, and cell death(9). Among them, cell adhesion is the ability of a single cell to attach to another cell or extracellular matrix (ECM). The ECM is a highly dynamic structure present in all tissues that undergoes controlled remodeling. During this process, quantitative and qualitative changes of its components occur in order to control homeostasis and tissue architecture(10, 11).The ECM is composed of various glycoproteins such as collagens, laminins, and fibronectins, and there are dozens of cellular receptors that directly interact with ECM components, for example, integrins or cadherins(10, 12).The interaction between ECM and receptors on the cell surface regulates cell behavior and plays an important role in cell-to-cell communication, cell proliferation, adhesion, and migration(13).

Recent discoveries have shown the essential role of ECM in regulating inflammatory responses, including cell extravasation and uptake, immune cell differentiation, polarization, activation, and retention in tissues(14, 15).

Notably, cellular matrix proteins are highly expressed in developing tissues and show lower expression levels in healthy mature tissues(16). However, the expression of matricellular proteins is often increased in response to injury and/or stress, particularly at sites of ECM remodeling(17-20). Due to the increased expression of matricellular proteins in injuries and diseases, the concept that matricellular proteins influence the recruitment and activity of immune cells is increasingly accepted(18, 19, 21, 22). Indeed, a number of studies have supported the critical and distinct functions of different matricellular proteins in inflammation(14, 23). Advanced researches about inflamation interactions in ECM shed the light for recently attractive cascade named, Lectin signaling pathway(24, 25). Mannose-binding lectin (MBL), known as mannan-binding protein (MBP), is a protein that is involved in complement activation via the lectin pathway(25, 26). The complement system provides immediate defense against infection and has proinflammatory effects(26).Matrix metalloproteinases (MMPs) are another group of zinc-dependent ECM endopeptidases capable of degrading almost all ECM components(27).The DKC1 gene is a type of MMPs that provides instructions for making a protein called dyscrein(28). This protein plays a role in maintaining structures called telomeres that are found at the ends of chromosomes. Telomeres help protect chromosomes from abnormally sticking together or breaking(29). In this regard, in the biophysical part of ECM, there are two very important signaling pathways, including Hippo and JAK/STAT, which play a vital role in the inflammatory response(30-32).

HNF4α in Hippo pathway as downstream effectors of YAP/TAZ activation in the cell fate determination and as well as colitis plays a role(33, 34). In the intestine, genetic deletion of HNF4α results in loss of mucin-related genes, increased intestinal permeability, loss of intestinal stem cell renewal, and susceptibility to inflammatory bowel disease(35).

The STAT family consists of six members, numbered 1 to 6, which have important structural and functional properties. All family members contain a so-called SRC-homology 2 (SH2) domain, a motif that allows specific binding to phosphorylated tyrosine residues. These tyrosine residues are present in the cytoplasmic tail of tyrosine kinase receptors (such as c-fms colony-stimulating factor receptors or platelet-derived growth factor receptors) and cytokine receptors (such as interleukin and interferon receptors). (36) That STAT3 activation in CD4+ T cells promotes IBD. Uncertainty remains, however, regarding which cellular compartment(s) STAT3 is most critical for IBD development and whether STAT3 in aggregate across all cells promotes or prevents IBD(37). Many studies have shown that targeting STAT3 signaling reduces polyp growth in animals carrying constitutive mutations of SKT11, the gene that encodes the tumor suppressor liver kinase B1 (LKB1), which is found in patients with Putz-Jeggers syndrome. It has mutated. Development of gastrointestinal polyps that predispose to CRC(38, 39).

In this study, we investigated the expression of selected genes in the ECM substrate and the inflammatory pathways effective in it, in patients with IBD separately from UC and CD, and also compared them in the control group for early surgical prognosis of these patients.

## 2. Materials and methods

### 2.1 Tissue collecting samples

This descriptive analytical study analyzed 50 patients with UC and 50 patients with CD. The samples that were collected were from patients who underwent surgery. These patients were all suffering from advanced IBD and were referred to the surgical department of Taleghani Hospital for surgery. Two expert pathologists, blinded to the study objectives, confirmed the pathologic features of the samples at Taleghani Hospital. The study adhered to the ethical guidelines outlined in the Declaration of Helsinki, including obtaining written consent from all human research participants. All procedures were conducted under the oversight of the Ethics Committee of the Shahid Beheshti University of Medical Sciences in accordance with the university’s policies on medical and research ethics.

### 2.2 Reverse transcriptase PCR (RT-PCR)

Total RNAs were extracted from obtained samples (Yekta Tajhiz Azma kit, Cat No. YT9065, Teheran, Iran). The RNAs concentration was measured by Nanodrop (NanoDrop Technologies, Wilmington, DE, USA) for acceptable quantification and qualification. RNAs were converted to cDNA by Retro transcriptase (RT) reaction (Yekta Tajhiz Azma kit, Cat No. YT9065, Teheran, Iran). The following reagents were added into a sterile nuclease-free tube on ice by order as mentioned: 0.1 ng of justified total extracted RNA were picked up and mixed with 1.0*μ*l of Random hexamer primer, then DEPC-treated water was added up to 13.5*μ*l. The ads in mixed gently, and incubated at 70°C for 5 min as manufacturer protocol. Denaturation was performed at 95°C for next 5 min and the following cDNA Synthesis Mix according to protocol was mixed up. The ingredients were as follow: 4*μ*l of 5x first-strand buffer, 1*μ*l of dNTPs (10 mM each), 0.5*μ*l of Rnasin (40U/μl), and 1*μ*l M-MLV enzyme. The mixture was added to the denatured RNA and incubated for 60 min at 37°C. Finally, the reaction was terminated by heating at 70°C for 5 min. The synthesized cDNAs were kept at -20°C until use. Primers for the loading control GAPDH generated a PCR product of 105 bp. Gene-specific primers for MASP2, DKC1, HNF4A and STAT3 were designed to produce PCR products of 96, 134, 145 and 75 bp respectively.

Primer sequences were as follows (for MASP2, DKC1, HNF4A and STAT3 specificity conferred by the 3′ end of the primer) (Table1).

**Table 1.**
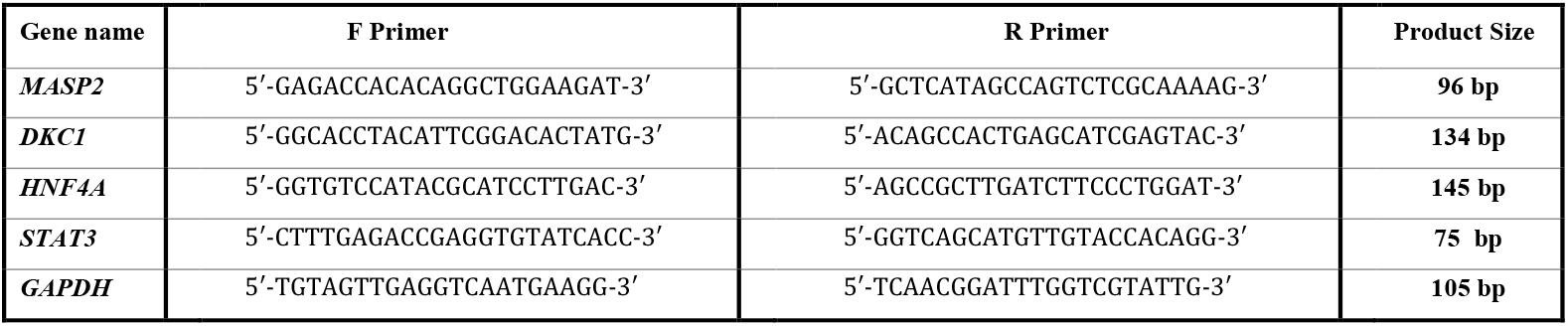
Sequences of primers used for Real-time RT-PCR (F forward, R reverse).

### 2.3 Real time-PCR

Following cDNA synthesis, MASP2, DKC1, HNF4A and STAT3 genes expression level was evaluated by Real-time PCR via SYBER Green qPCR method. The quantification was performed by use of MasterMix 2x (Yektatajhiz kit, Cat No YT2551, Tehran, Iran). Real-time PCR was carried out by designed primers as mentioned in (Table 1). The justification of the primers was estimated by Linreg method and the efficiency of each primer was calculated 95%. The Real time reactions were administered under the following condition: pre-denaturation at 95°C for 20 s in one cycle, denaturation at 95°C for 10 s, annealing on 60°C for 10 s, extension on 72°C for 20 s, 40 cycles repeat all three main levels. Amplification signals were normalized by glyceraldehyde 3-phosphate dehydrogenase (GAPDH) as house-keeping gene. The relative quantification and fold changes were evaluated by the 2^−ΔΔCt^ method.

### 2.4 Real time-PCR Statistical analysis

All released RT-PCR data were analyzed by GraphPad Prism version 9 software (San Diego, CA, USA). The data were not-normally distributed and the non parametric tests were applied. The Mann Whitney and between-groups comparisons were tested for statistical significance using the paired t-test and one-way analysis of variance (ANOVA), respectively. All P-values < 0.05 were considered statistically significant.

### 2.5 Adhesion Molecule Genes Correlation in Healthy Tissue

To investigate the co-expression of MASP2, DKC1, HNF4A and STAT3 genes at the transcript level healthy colon tissue, the “correlation analysis” tool from the GEPIA2 database (http://gepia2.cancer-pku.cn) and the Pearson correlation coefficient method were utilized. The correlation of gene expression was evaluated one by one in sigmoid and transverse colon samples.

### 2.6 Adhesion Molecules Co-expressed genes and PPI Network

Protein-protein interaction (PPI) network of MASP2, DKC1, HNF4A and STAT3 was displayed by the STRING database (http://string-db.org). The minimum required interaction score was set at 0.4. Then, nodes with edges < 1 and nodes not involved in the leading integrated network were excluded.

## 3. Results

### 3.1 Clinicopathological Characteristics of Patients with UC & CD

Clinicopathological features of the 50 included patients with UC and 50 included patients with CD were extracted from their files and presented at the beginning of the survey. The mean age of the patients was (60.32 ± 10.17 years). The analyzed data was presented in Table 2 & 3. The 23-years-old woman was the youngest and the 61 years-old man was the oldest patient who entered the study. Although 75% of patients with IBD verified positive family history of colon diseases but no significant P-value was admitted to familial history. In present survey, tissue detected in distal-left colon (including the rectum, sigmoid colon, descending colon) were significantly (P-value < 0.022) classified in UC category. Also tissue detected in proximal-right colon (including the cecum, ascending colon, transverse colon) were significantly (P-value < 0.022) classified in CD category (Table 2, 3).

**Table 2.**
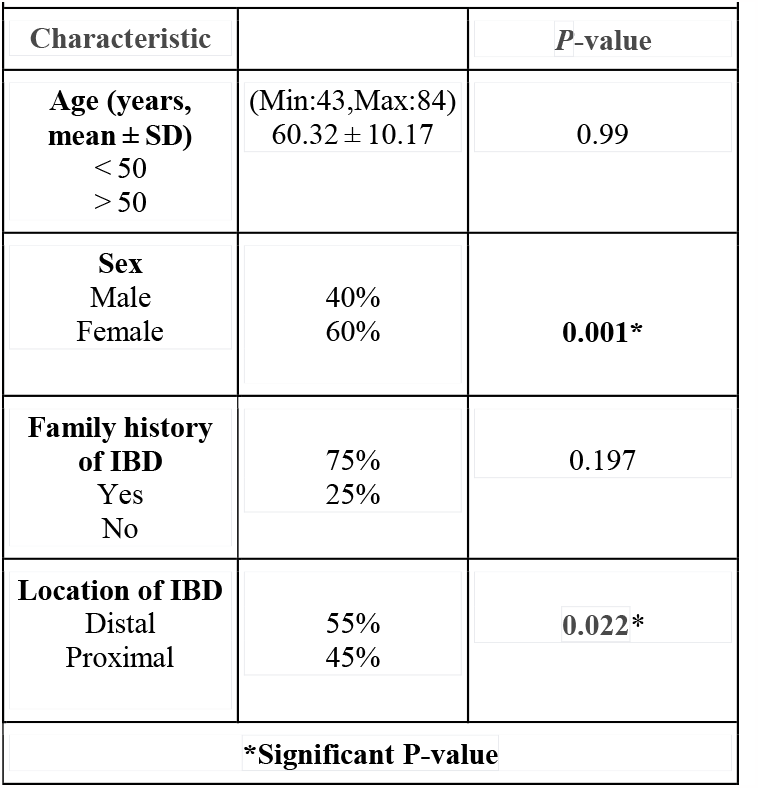
Descriptive analysis of clinicopathological characteristics based on the histology of IBD (UC).

| Characteristic |  | P-value |
| --- | --- | --- |
| **Age (years, mean ± SD)**<br>< 50<br>> 50 | (Min:43,Max:84)<br>$60.32 \pm 10.17$ | 0.99 |
| **Sex**<br>Male<br>Female | 40%<br>60% | **0.001\*** |
| **Family history of IBD**<br>Yes<br>No | 75%<br>25% | 0.197 |
| **Location of IBD**<br>Distal<br>Proximal | 55%<br>45% | **0.022\*** |

\*Significant P-value

**Table 3.** Descriptive analysis of clinicopathological characteristics based on the histology of IBD (CD).

| Characteristic |  | P-value |
| --- | --- | --- |
| **Age (years, mean ± SD)**<br>< 50<br>> 50 | (Min:43,Max:84)<br>$60.32 \pm 10.17$ | 0.99 |
| **Sex**<br>Male<br>Female | 40%<br>60% | **0.000\*** |
| **Family history of IBD**<br>Yes<br>No | 88.9%<br>9.1% | **0.001\*** |
| **Location of IBD**<br>Proximal<br>Distal | 55%<br>45% | **0.022\*** |

\*Significant P-value

### 3.2 Selected Cell Adhesion Genes in IBD (UC & CD)

The mRNA expression of desired genes was measured by Real-time PCR technique and the RQ of each gene was calculated via 2^−ΔΔCt^ method. The expression analyzed data were subdivided to over and under RQ = 1 to represent up and down regulation of genes.

The confirmed significant clinicopathological parameters were evaluated with nominated genes expression analysis. The MASP2 and DKC1 were significantly related with UC. Down regulation of MASP2 in Proximal colon with 33.3% of cases and up regulation in distal with 75.7% was seen in SPSS analysis with P-value < 0.017. More than 72% of cases with upregulated DKC1 was seen in distal colon and only 27% of UC in distal was down regulated in DKC1. In the samples related to CD as well the up regulation of HNF4A and STAT3 were significantly. Down regulation of HNF4A in Distal colon with 36.2% of cases and up regulation in Proximal with 81.7% was seen in SPSS analysis with P-value <0.019. (Table 4).

**Table 4.** Descriptive analysis of clinicopathological characteristics based on the histology of IBD (UC & CD).

| Parameters: | Tissue |  |  | Location |  |  |
| --- | --- | --- | --- | --- | --- | --- |
|  | UC | Normal | P-value | Distal | Proximal | P-value |
| **Genes:** |  |  |  |  |  |  |
| *MASP2* | 88.9% | 39.3% | **0.010\*** | 75.7% | 33.3% | **0.017\*** |
| *DKC1* | 88.9% | 45.5% | **0.03\*** | 72.7% | 27.3% | 0.64 |
| Parameters: | Tissue |  |  | Location |  |  |
|  | CD | Normal | P-value | Distal | Proximal | P-value |
| **Genes:** |  |  |  |  |  |  |
| *HNF4A* | 88.9% | 39.3% | **0.010\*** | 75.7% | 33.3% | **0.019\*** |
| *STAT3* | 75.6% | 44.7% | **0.02\*** | 81.7% | 52.4% | 0.61 |

### 3.3 Expression Level of Genes in IBD (UC & CD)

Comparing the expression level of MASP2 and DKC1 based on the UC and normal samples showed a diminished expression level of MASP2 in normal in contrast with UC cases (P-value: 0.0057). The expression level difference between UC and normal categorized samples type in DKC1 was near significant with up regulation preference in UC (P-value: 0.053). In cases of CD also expression level of HNF4A and STAT3 admitted the significant up regulation of CD samples in contrast with normal expression levels showed substantial differences between IBD and normal samples (P-value: 0.027) (Figure 1).

**Figure 1.**
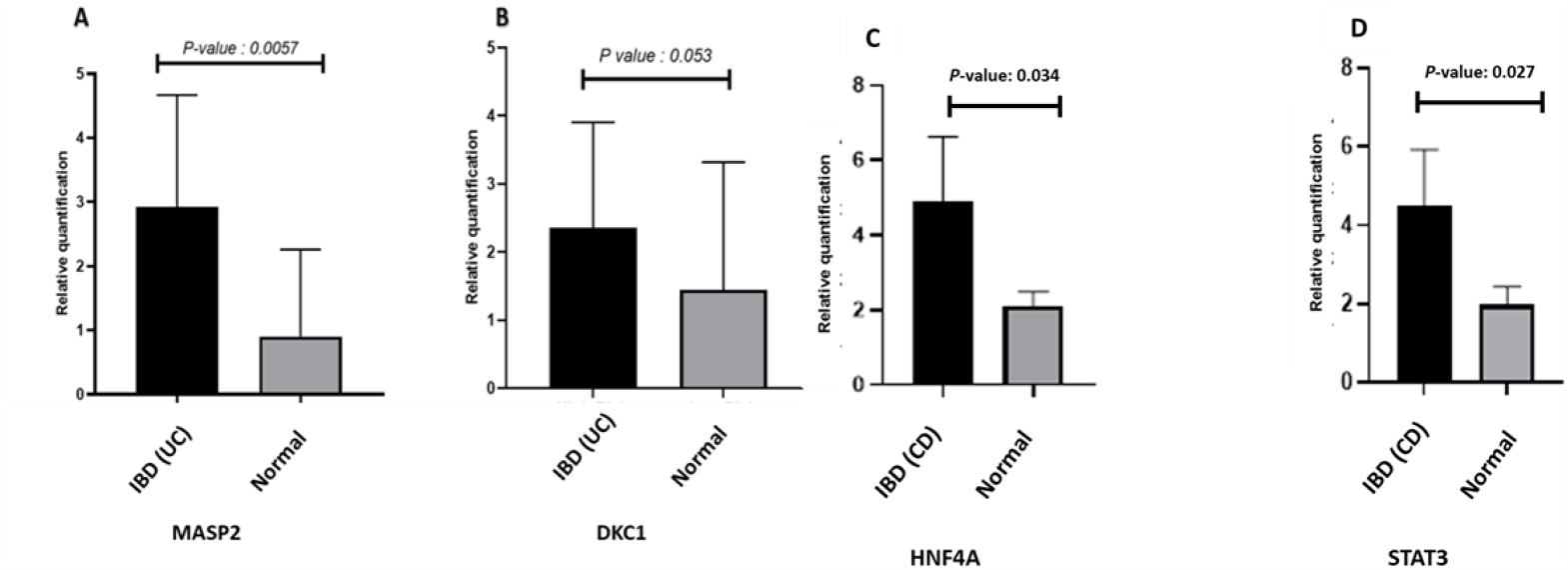
Expression levels of MASP2, DKC1, HNF4A and STAT3 in IBD (UC & CD).

### 3.4 Prognostic Biomarker Determination In advance IBD (UC & CD)

Additionally, a receiver operating characteristic (ROC) curve was plotted to assess the prognostic value of MASP2, DKC1, HNF4A and STAT3 expression in patients with IBD. The differentiation between advance IBD samples and normal samples determined by criteria mentioned above. The AUC for MASP2 was evaluated 0.87 by sensitivity of 100% and specificity of 78% (P-value = 0.0071). The DKC1 and HNF4A expressions with 77% area under the curve demonstrates sensitivity of 100% and specificity of 67% with significant P-value to detect between advance IBD and normal samples of colon and rectum. Also the estimated area for STAT3 ROC was 0.80 with sensitivity of 67% and specificity of 90% (P-value = 0.027). (Figure. 2)

**Figure 2.**
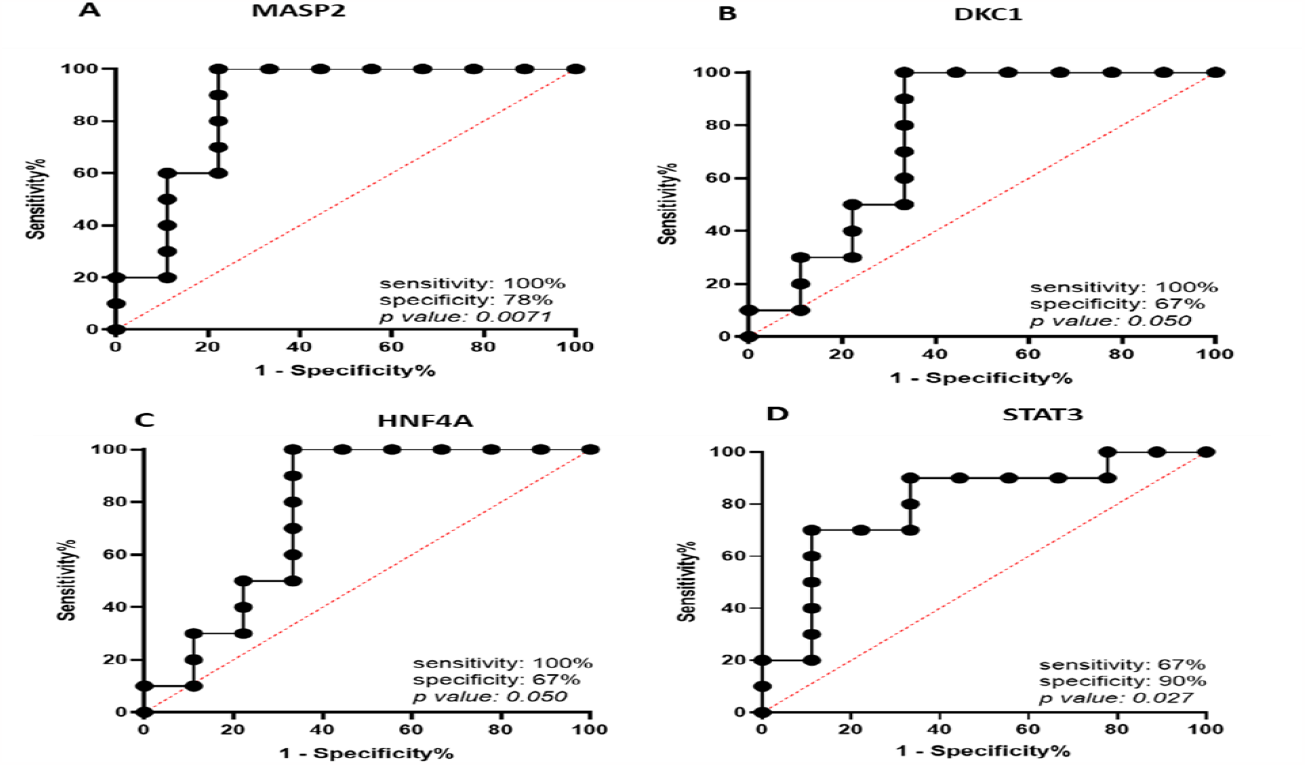
ROC Curves Showing AUC, Sensitivity, and Specificity of Selected Genes as Biological Biomarkers

### 3.5 Correlation of MASP2, DKC1, HNF4A and STAT3 Genes Expression in Healthy Colon

To validate the co-expression of of MASP2, DKC1, HNF4A and STAT3 at the transcript level in healthy sigmoid and transverse colon data from GTEx, the “correlation analysis” tool based on RNA-seq data and Pearson’s correlation coefficient from the GEPIA2 database was executed. According to the obtained results, the highest correlation was seen between MASP2 and DKC1 & HNF4A and STAT3 genes. Based on this, MASP2 and DKC1 show a positive and significant correlation at the transcript level (P-value= 0.024 and R=0.17). In the same direction of the correlation between the expression of HNF4A and STAT3 genes was significant (P-value= 0.0038 and R=0.16) (Figure 3).

**Figure 3.**
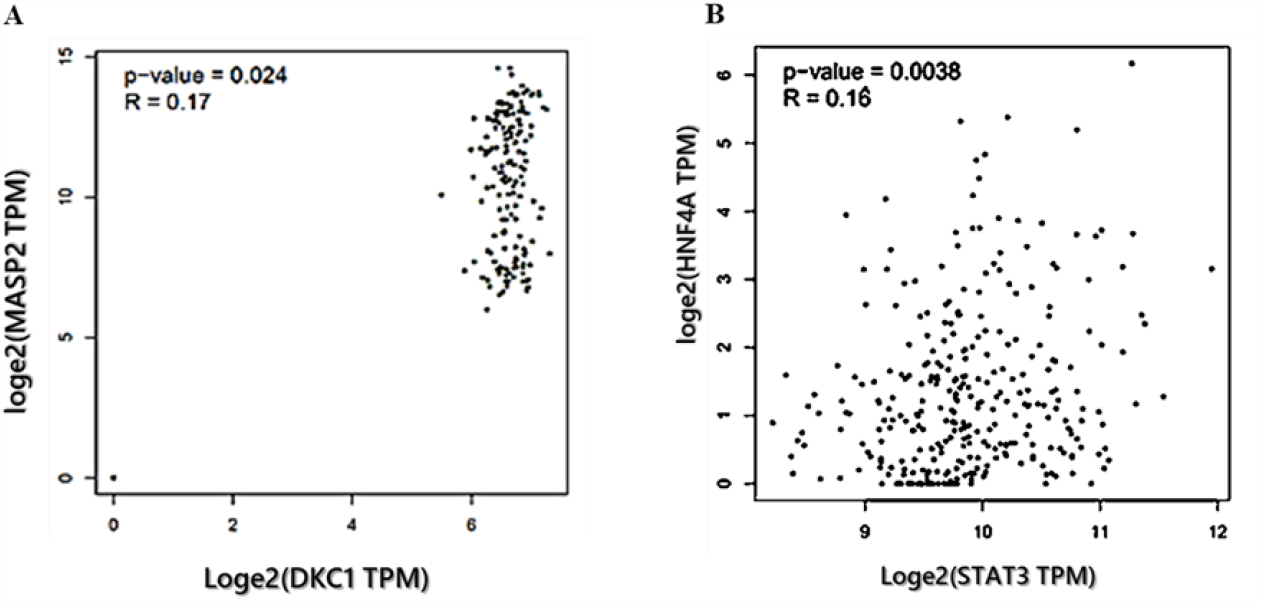
Correlation of *MASP2 and DKC1 & HNF4A and STAT3* genes expression. **(A)** The correlation of MASP2 and DKC1 at transcriptome level in GTEx transverse and Sigmoid colon. **(B)** The correlation of HNF4A and STAT3 at transcriptome level in GTEx transverse and Sigmoid colon.

### 3.6 PPI network of Adhesion Molecule Genes

We also retrieved the PPI network, including the top proteins associated with *MASP2 and DKC1 & HNF4A and STAT3* genes from STRING, which were supported by experimental data. shows the interaction network of the mentioned genes and proteins. Furthermore, in order to identify common proteins among these adhesion prominent genes and to find out the expected functional mechanism of selected genes, the proteins present in the PPI network of genes were intersected, which is displayed in (Figure 4. A & B). Intersection analysis of PPI networks between MASP2 and DKC1 illustrated those six proteins, including TERT, GAR1, FBL, MBL2, NOP10, and KAT5.This is while eight common proteins were found among the PPI network of the HNF4A and STAT3 explored genes, these proteins including: NANOG, SRC, EGFR, CDC37, STUB1, JAK1, JAK2 and NR3C1(Figure 4. C & D).

**Figure 4.**
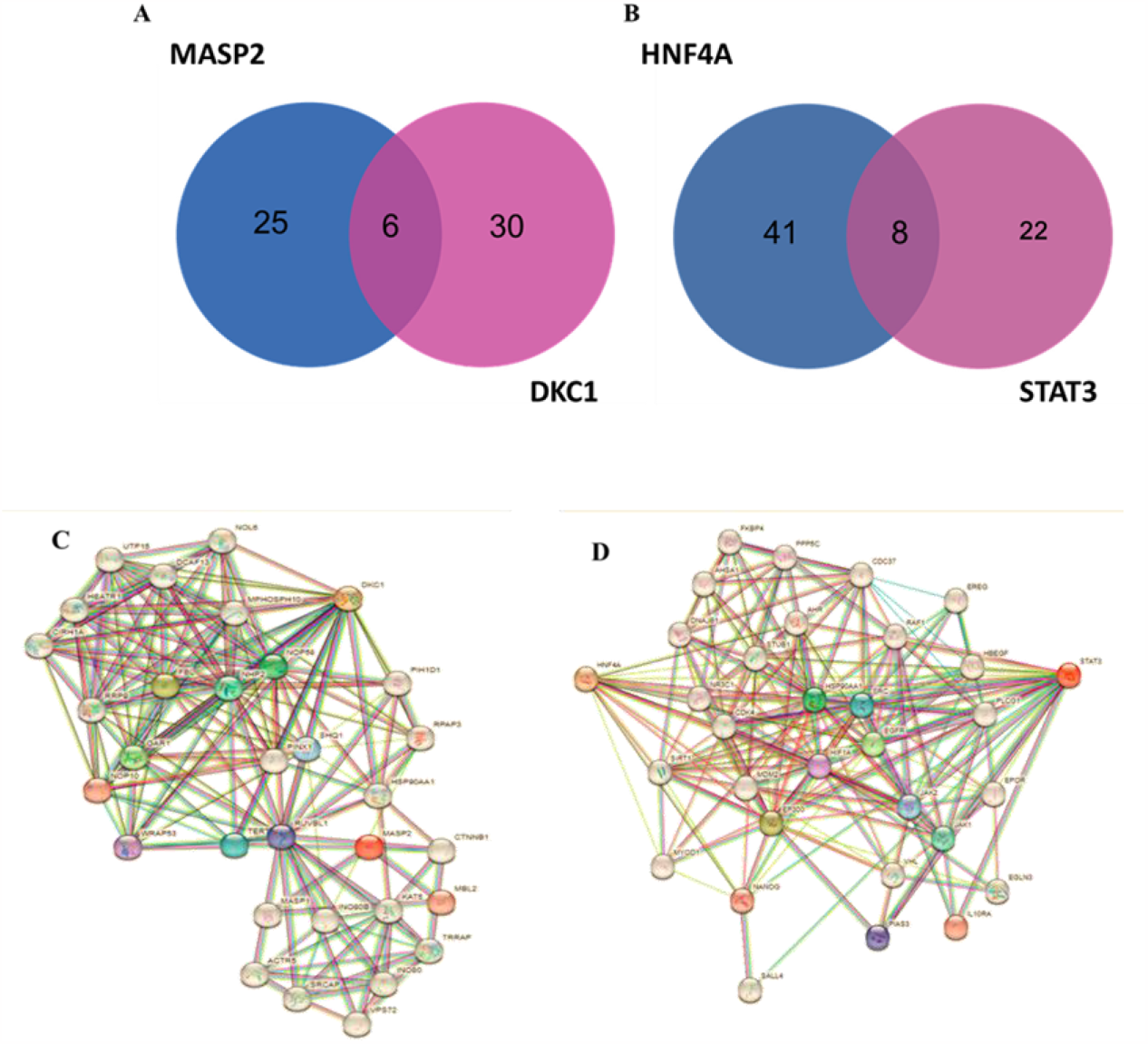
PPI network of *MASP2 and DKC1 & HNF4A and STAT3* genes. **(A)** PPI network of 40 MASP2 and DKC1 -binding proteins. **(B)** PPI network of 40 HNF4A and STAT3 -binding proteins. **(C)** Intersection analysis of proteins present in *MASP2* and *DKC1* genes PPI networks. **(D)** Intersection analysis of proteins present in *HNF4A and STAT3* genes PPI networks.

## 4. Discussion

Ulcerative colitis (UC), a subtype of inflammatory bowel disease (IBD), is a chronic relapsing and progressive disorder with unclear etiology (40). Current therapeutic drugs for UC include 5-aminosalicylic acid (5-ASA) drugs, steroids, immunomodulators, and biological agents, and promising therapeutic drugs targeting specific signaling pathways involved in the pathogenesis of IBD have continued to be investigated in recent years(41). For some patients, 5-ASA therapy is sufficient for clinical remission induction and maintenance during the relatively long-term course after diagnosis, while others require strict interventions at diagnosis or need treatment escalation within a shorter time(42). Accordingly, reliable prediction for the disease course of patients with UC is of great significance to guide therapeutic decision-making and follow-up management, but it is still a challenge.

In addition to these cases in terms of mental problems, IBD disease has an undoubted impact on patients’ quality of life with important consequences at psychosocial level (43, 44). However, psychosocial support for IBD care is still underestimated and there is no clear consensus on priorities for interventions. In addition, although patient engagement is a crucial target in chronic care, it often appears to be an abstract principle and it is not applied to clinical practice (45).

Advances in our understanding of immune and non-immune cell interactions with the ECM in IBD have revealed several previously unsuspected mechanisms involved in disease pathogenesis. Previously, ECM changes in IBD were regarded as a consequence of inflammation, but emerging evidence suggests that ECM remodeling (destruction and de novo synthesis) is an active participant in the inflammatory process in the pathobiology of IBD (46).

In previous studies, intensification therapy in medicine and the requirement for surgery were generally considered as signs of complicated or aggressive disease course in patients with UC. To provide more effective therapy approaches for these patients at the appropriate time, researchers have conducted multiple studies to find disease-course-associated predictors at diagnosis. However, few data are available on the indolent course of UC and its related factors (8). Recently, Yanai et al. defined indolent course in Crohns disease as a disease course without the need for strict interventions (including steroids, immunomodulators, biological agents, hospitalization, or surgery therapy) during the whole follow-up period. As noted above, we believe that the definition of indolent course in Crohns disease may also apply to patients with UC (47). Considering the prevalence of this autoimmune disease, especially in developing countries, and also considering the lowering of the age at which it is diagnosed and observed in young people, the study of genes and non-coding RNAs related to them in inflammatory molecular pathways is very worthwhile. Attention and prognosticator.

In this regard, in this study, an attempt has been made to introduce the panel of markers involved in IBD disease by separating it into two related types (CD, UC); So that we can use them to perform surgery on patients who need surgery without wasting time and money and reduce the psychological burden of the patient. Considering the existing concerns in IBD patients and also to avoid wasting time to treat these patients, especially in teenagers, this study investigates and presents a gene panel separating both types of UC and CD Involved in ECM include.

We studied HNF4A and STAT3 in CD, as well as MASP2 and DKC1 in UC patients; Based on this group of genes involved in inflammatory pathways, we can identify and evaluate patients who need early surgery.

## Limitation of Biomarkers

Ideal biomarkers should be non-invasive, sensitive, disease-specific, easy to implement, and cost effective.

## Ethics approval

This study was approved by the Ethics Committee of Shahid Beheshti University of Medical Sciences, Tehran, Iran with the registration number of: IR.SBMU.MSP.REC.1400.623.

## Informed consent

Informed Consent Written consent was obtained from all patients who were informed that the data would be used for research. Also the use of human specimens is also based on Declaration of Helsinki.

